# Secondary neurulation-fated cells in the tail bud undergo self-renewal and tubulogenesis regulated by a Sox2 gradient

**DOI:** 10.1101/295709

**Authors:** Teruaki Kawachi, Eisuke Shimokita, Ryosuke Tadokoro, Yoshiko Takahashi

**Affiliations:** Department of Zoology, Graduate School of Science, Kyoto University Kitashirakawa, Sakyo-ku, Kyoto; Graduate School of Biological Sciences, Nara Institute of Science and Technology, 8916-5, Takayama, Ikoma, Nara, 630-0192, Japan; AMED Core Research for Evolutional Science and Technology (AMED-CREST), Japan Agency for Medical Research and Development (AMED), Chiyoda-ku, Tokyo 100-0004, Japan; Present address: Department of Anatomy and Cell Biology, Institute of Biomedical Sciences Tokushima University Graduate School, 3-18-15 Kuramoto-cho, Tokushima 770-8503, Japan

## Abstract

During amniote development, anterior and posterior components of the neural tube form by primary neurulation (PN) and secondary neurulation (SN), respectively. Unlike PN, SN proceeds by the mesenchymal-to-epithelial transition of SN precursors in the tail bud, a critical structure for the axial elongation. Our direct cell labeling delineates non-overlapping territories of SN- and mesodermal precursors in the chicken tail bud. SN-fated precursors are further divided into self-renewing and differentiating cells, a decision regulated by graded expression levels of Sox2. Whereas Sox2 is confined to SN precursors, Brachyury (T) is widely and uniformly distributed in the tail bud, indicating that Sox2^+^/Brachyury^+^ cells are neural-fated and not mesodermal. These results uncover multiple steps during the neural posterior elongation, including precocious segregation of SN precursors, their self-renewal, and regulation by graded Sox 2.

## Introduction

In amniotes (mammals, birds and reptiles), the neural tube is composed of a neuroepithelium that forms through two processes along the anterior posterior (AP) axis. These processes are primary neurulation (PN) and secondary neurulation (SN). The PN occurs in the anterior region of the body comprising the future brain and spinal cord in thoracic regions, where an epithelial cell sheet (neural plate) invaginates to make a tubular structure (Colas & Schoenwolf, 2001, Copp *et al.*, 2003, Saitsu *et al.*, 2004). In contrast, the SN-mediated neural tube, located posterior to the lumbar/hind limbs (in chickens and humans), is achieved by the mesenchymal-to-epithelial transition (MET) of SN precursors in the tail bud (Hughes & Freeman, 1974, Pasteels, 1937, Schoenwolf, 1979, Holmdahl, 1938). The tail bud is a transient structure located at the posterior end of the embryo during axial elongation, and it continuously supplies cells to the region of the neural tube undergoing SN (designated as the secondary neural tube). The secondary neural tube eventually provides innervations that govern physiological functions for a wide variety of organs including the colon, bladder, genitalia, and tail movement (Copp *et al.*, 2015, Le Douarin & Kalcheim, 1999, Snell, 1995, Stiefel *et al.*, 2007). In addition, a failure of SN is implicated to be a cause of neural tube defects (NTDs) including *spina bifida*, which are among the most common congenital malformations in humans (Copp *et al.*, 2015, Copp *et al.*, 2013, Dady *et al.*, 2014, Detrait *et al.*, 2005, Greenberg *et al.*, 2011, Marks & Khoshnood, 1998). It is therefore important to elucidate the mechanisms by which the secondary neural tube formation is regulated. However, the ways that the SN precursors accomplish the tasks of neural differentiation and tubular morphogenesis remain poorly understood.

The tail bud also provides cells that form the paraxial mesoderm of the posterior embryo, as well as the midline components of the notochord and floor plate during the axial elongation (Cambray & Wilson, 2002, Catala *et al.*, 1996, Catala *et al.*, 1995). However, how these different lineages become specifically located in the tail bud remains undetermined, although several different hypotheses have been proposed. A main reason for our ignorance is that cells constituting the tail bud are mesenchymal in shape, hampering morphological delineation between the lineages (Colas & Schoenwolf, 2001, McGrew *et al.*, 2008). To solve this long-standing question, a cell-labeling technique that directly visualizes SN precursors must be useful.

We previously used direct cell labeling in the embryonic chicken tail bud (Shimokita & Takahashi, 2011) to show that SN-precursors are derived from a specific region of the epiblast, called the presumptive SN (preSN). This preSN is located along the midline posterior to the Hensen’s node at Hamburger and Hamilton stage 8 (HH 8; Fig. 1A-D). The preSN at this stage is unique for two features. First, this specific region of epiblast is not underlain by the primitive streak. Rather, the anterior end of the primitive streak is apart posteriorly from the preSN by 200-250 µm (Fig. 1A). Second, the preSN lacks the underlying basement membrane, thereby allowing preSN-epithelial cells to ingress by the epithelial-to-mesenchymal transition (EMT) in a manner similar to mesodermal ingression from the primitive streak. Labeling either by DiI or by EGFP-electroporation revealed that the preSN-derived ingressed cells constitute the tail bud where they remain mesenchymal until they are incorporated into the secondary neural tube (Shimokita & Takahashi, 2011). Little or no mesodermal cells are derived from the preSN region.

**Figure 1.**
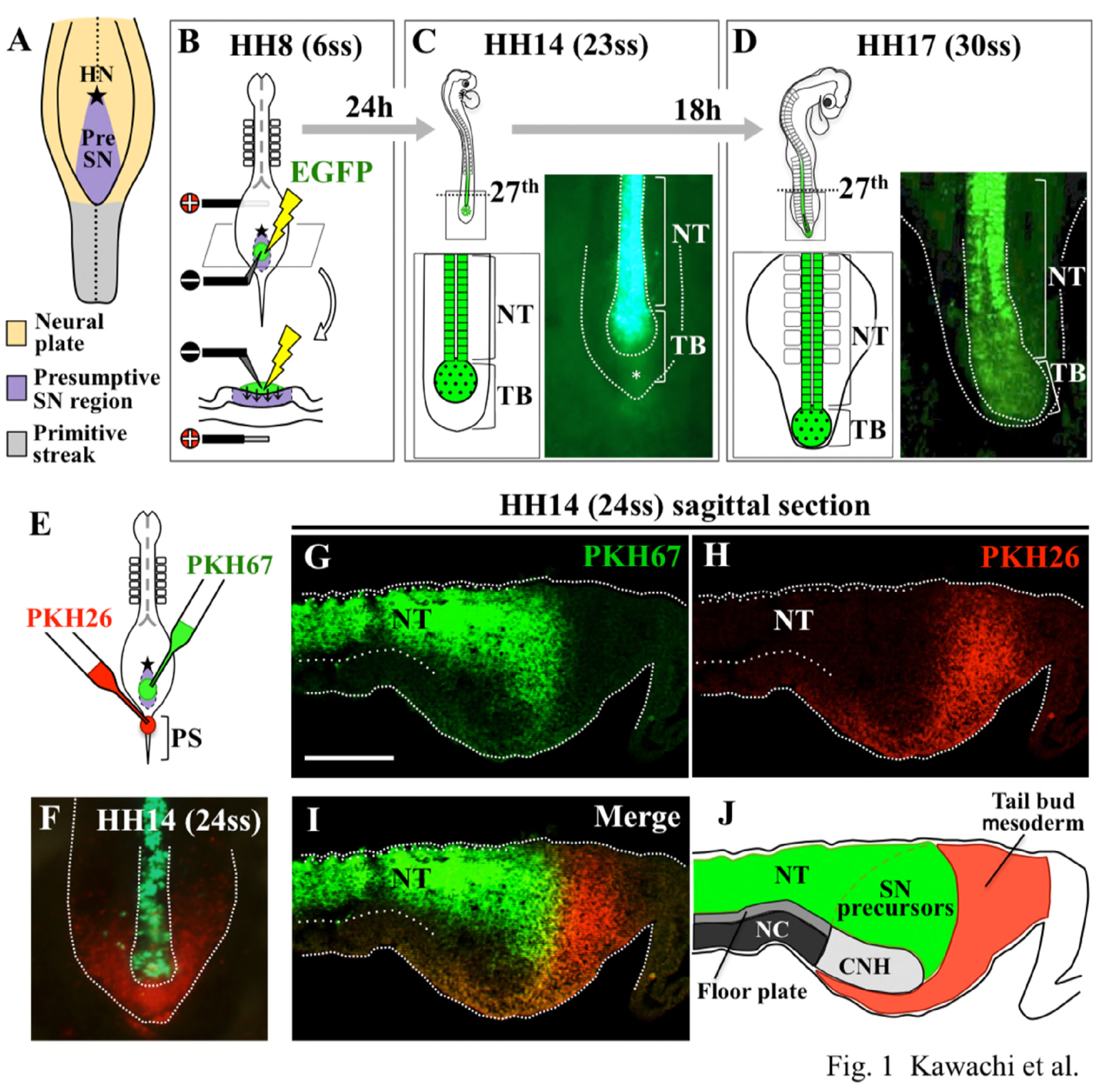
Segregation of SN- and mesoderm-fated cells in the tail bud at HH14. (A) A fate map of the HH8 epiblast caudal to the Hensen’s node (HN) modified from Shimokita and Takahashi (2011). A specific region called presumptive secondary neurulation (preSN, purple) gives rise solely to secondary neurulation (SN), which undergoes posteriorly to 27^th^ somite level. The EGFP-positive neural tube anterior to 27^th^ is of primary neurulation. (B) *In ovo* electroporation conducted at HH8 (see also Materials and Methods). (C, D) EGFP-electroporated preSN gave rise to EGFP-positive cells in the SN-forming cells but not in the mesoderm as seen at HH14 and HH17. (E) Simultaneous labeling by different colors of PKHs of preSN and anterior primitive streak (mesoderm) in a single embryo. (F) Dorsal view, and (G-I) sagittal sections at HH14. Little overlapping between SN (green)- and mesoderm (red)-fated cells in the tail bud. CNH, notochord and floor plate were unlabeled. (J) Schematic trace of SN- and mesodermal precursors in the tail bud. NT, neural tube; TB, tail bud; CNH, chord neural hinge; NC, notochord, *, background. Scale bar; 100 µm.

By extending these findings, we noticed that despite the massive elongation of the secondary neural tube that continuously receives SN precursors, the amount of labeled SN precursors in the tail bud remains constant as seen at HH14 (23ss) and HH17 (30ss) (Fig. 1B-D). This raised the possibility that SN precursors would possess a self-renewing ability in the tail bud. In this study, we have tested this possibility by following a precise delineation of the SN-fated cell population in the tail bud. We have found that the SN-precursors in the tail bud are composed of two groups of cells, self-renewing cells and neural-specified transitional cells. The SN precursor territory including these two subpopulations expresses both Sox 2 and Brachyury, with the Sox 2 level decreasing posteriorly. This Sox 2 gradient plays roles in the regulation of SN precursors in the tail bud.

## Material and Methods

### Chicken embryos

Fertilized chicken eggs were obtained from Shiroyama poultry farm (Sagamihara, Japan). Embryos were staged according to Hamburger and Hamilton (Hamburger & Hamilton, 1951) or the somite number. All animal experiments were conducted with the ethical approval by Kyoto University (No. H2620).

### Vector constructions

pCAGGS-EGFP was as previously described (Momose *et al.*, 1999). cDNAs encoding Sox2 and Sox3HMG-EnR were provided by Drs. K. Fukuda (Tokyo Metropolitan University) and Y. Sasai (CDB Riken), respectively. The cDNAs were individually subcloned into the Mul I-Nhe I site of pBI-EGFP or pBI-DsRed vector (Watanabe *et al.*, 2007).

### *In ovo* electroporation

An anode and cathode were prepared with a platinum wire (diameter of 0.3-0.5 mm) and a sharpened tungsten needle (40 µm diameter at the tip), respectively. The *in ovo* electroporation was carried out at HH8 according to the method previously reported (Momose *et al.*, 1999, Nakaya *et al.*, 2004) with slight modifications: the anode was inserted in between the embryo and yolk, and a DNA solution containing 2% fast green FCF (Nakarai) was laid on the epiblast, followed by electric charges five times of 5 V, 25 ms with 100 ms intervals (Electro Square Porator ECM830; BTX) using the cathode. Maximal attention was paid so that the DNA solution did not spread into the primitive streak, from which mesoderm arises.

### PKH-labeling

The presumptive SN region in the epiblast or primitive streak of HH 8 (6 somites stage) embryos were labeled with PKH26 Red Fluorescent Cell Linker (Sigma) or PKH67 Green Fluorescent Cell Linker (Sigma) using a micropipette pulled from a 1 mm glass capillary in a vertical micropipette puller (model PC-10; Narishige).

### *In situ* hybridization

Whole-mount in situ hybridization was performed as previously described (Henrique *et al.*, 1995, Atsuta & Takahashi, 2015) with some modifications. Embryos were fixed overnight in phosphate-buffered saline, PBS (0.1M Tris-HCl [pH 7.5], 0.15M NaCl) containing 4% paraformaldehyde (PFA), 100 mmol ⁄ L 3-(N-morpholino) propanesulfonic acid (MOPS) (pH 7.4), 2 mmol ⁄ L ethylene glycol tetraacetic acid (EGTA), and 1 mmol⁄ L MgSO_4_ ⁄7H_2_O. After washing twice in 0.1 % Tween 20 in PBS (PBST), specimens were dehydrated by a successively graduated series of methanol in PBST (40%–100%). They were subsequently treated with proteinase K (20 µg ⁄ mL) for 15 min, followed by refixation in 4% PFA in PBS for 20 min at room temperature (RT). After two PBST washes, the embryos were transferred to hybridization buffer (ULTRAhyb, Ambion) and prehybridized for 1 hr at 68 °C. Hybridization was carried out overnight at 68 °C in the hybridization buffer containing a digoxigenin (DIG)-labeled RNA probe. The embryos were washed three times for 30 min each in 50% formamide, 5 x standard saline citrate (SSC), 1% ethylenediaminetetraacetic acid (EDTA), 0.2% Tween 20, 0.5% 3-[(3-Cholamidopropyl) dimethylammonio] propanesulfonate (CHAPS), at 68 °C. They were further washed in 0.1 mol ⁄ L Maleic Acid (pH 7.4), 0.15 mol ⁄ L NaCl, 1% Tween 20 (MABT) at least three times, prior to preblocking in 2% BBR in MABT. Hybridization was carried out by incubating embryos in blocking solution, which contained alkaline phosphatase-conjugated anti-DIG antibody (Roche), overnight. After the embryos were extensively washed in MABT for at least 5 h with several changes of solutions, they were processed to NTMT (100 mmol ⁄ L Tris-HCl [pH 9.5], 100 mmol /L NaCl, 50 mmol ⁄ L MgCl_2_, 0.1% Tween 20). The alkaline phosphatase activity was visualized by incubating embryos in NTMT containing 0.45 mg⁄mL nitroblue-tetrazolium chloride (Roche) and 0.175 mg⁄mL 5-bromo-4-chloro-3-indolyl phosphatase (Roche). After stopping the color reaction, embryos were post-fixed in 0.1% glutaraldehyde /4% PFA in PBS for 30 min at 4 °C.

### Immunohistochemistry

Embryos were fixed for 2 hours in PBS containing 3 % PFA at 4**°**C, followed by preparation of cryostat sections 10 µm thick. The detection of the Sox2 or Brachyury proteins (T) was performed as follows: after pre-blocking with 2 % blocking reagent (Roche)/PBS for 1 hr at RT, the sections were incubated at 4 °C overnight with either of the following antibodies in 1% blocking reagent (BBR, Roche) **⁄** PBS: anti-Sox2 mouse polyclonal antibody (ab5603; Millipore) (Oginuma *et al.*, 2017) diluted 1:100, and anti-Brachyury goat polyclonal antibody (AF2018; R&D Systems) (Nakanoh *et al.*, 2017) diluted 1:100, anti-Fibronectin antibody (F3648; Sigma) (Yoshino *et al.*, 2014), anti-GFP antibody (11814460001, Roche) diluted 1:300. After washing three times in PBST, the specimens were reacted with either of the following second antibodies: anti-rabbit IgG-Alexa 568 conjugated donkey antibody (A10042, Invitrogen), anti-goat IgG-Alexa 488 conjugated donkey antibody (A11055, Invitrogen), anti-rabbit IgG-Alexa 647 conjugated donkey antibody (A31573, Invitrogen), anti-mouse IgG-Alexa 488 conjugated donkey antibody (R37114, Invitrogen) diluted 1:400 with 1% blocking reagent **/**PBS for 1 hr at RT. The reaction was terminated by washing three times in PBS, and the sections were sealed by VectaShield (Funakoshi).

## Results

### SN precursors are segregated from mesodermal cells in the tail bud at HH14

To precisely locate the SN precursors and other lineages in the tail bud, we labeled the preSN region by PKH 67 (green) and the anterior primitive streak by PKH26 (red) in a single HH8 embryo having 6-somite (6ss). Dorsal views of these embryos at HH14 (24ss) showed a domain of preSN-derived cells that was distinct from that of anterior primitive streak-derived paraxial mesodermal precursors in the tail bud (Fig. 1A-D, n=123; Fig. 1E, F, n=26). The preSN-derived cells remained restricted to the secondary neural tube-forming territory with little, if any, contribution to the mesoderm. In sagittal histological sections, the reciprocal segregation is also obvious between the SN- and mesodermal populations in the tail bud: SN precursors (green) occupy the anterior half of the tail bud with the posterior half being populated by mesodermal cells (red) (Fig. 1G-J, n=12). As expected, neither SN-nor paraxial mesodermal precursors were found in the chord neural hinge (CNH), known to give rise specifically to the notochord and floor plate (Catala *et al.*, 1995). These observations demonstrate that a vast majority of cells of the tail bud are differentially fated at least at HH14. The segregation between SN- and mesoderm territories in the tail bud was retained at HH17 (30ss), where mesodermal population was located more ventrally to the SN precursors, and was not overtly visible in the dorsal view (Fig. 1D; n=4).

### SN precursors in the tail bud contain self-renewing cells

Since SN precursors are constantly present in the tail bud during the secondary neural tube elongation, we reasoned that these precursors contain self-renewing cells. To test this, we prepared HH17 (30ss) embryos in which SN precursors had been EGFP-labeled (Fig. 2A). From these EGFP-embryos, a piece was dissected from the SN precursor domain of the tail bud, and transplanted into the corresponding region of the tail bud at the presumptive 30-32 somite level of a non-electroporated HH14 embryo (24ss) (Fig. 2A-D; n=8). Even though the experimental approaches in this study allowed us to specifically trace the EGFP-labeled SN-precursors, we paid maximal attention not to take contamination of unlabeled ventral cells in the tail bud that contain CNH and some paraxial mesoderm-fated cells.

**Figure 2.**
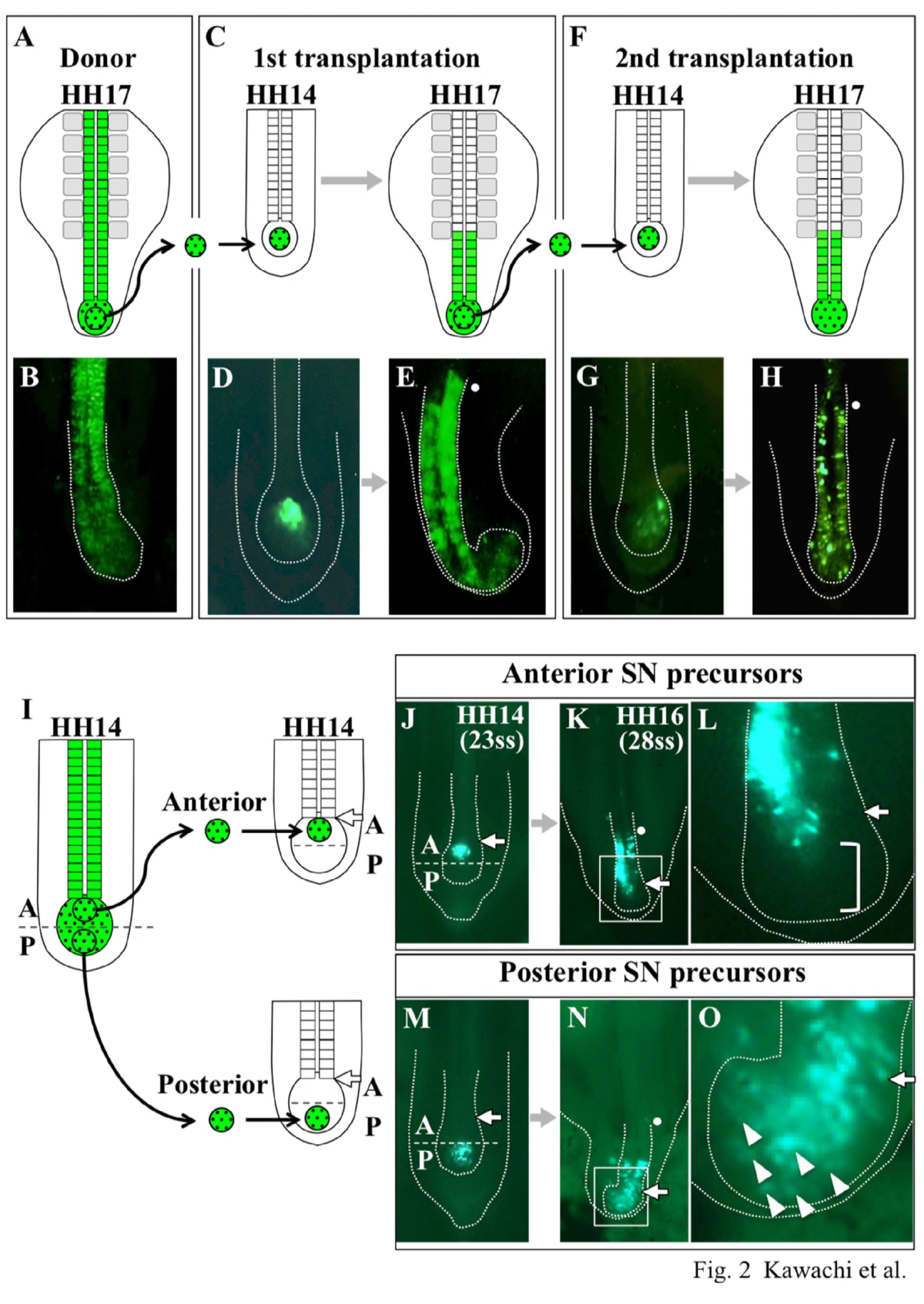
Transplantation with SN-fated cells revealed self-renewing cells in the posterior half of the SN-territory. (A-E) EGFP-expressing SN-fated cells were dissected from the tail bud of HH17 embryo, and transplanted into a corresponding region of the tail bud of HH14 embryos. (F-H) When the host embryo developed to HH17, similar transplantation was repeated. (I) The EGFP-positive SN precursor territory in the tail bud was subdivided into the anterior and posterior halves, and each of them was isotopically transplanted into a host embryo (HH14). (J-L) The anterior SN precursors underwent neural tube differentiation without remaining in the tail bud (bracket). (M-O) The posterior SN precursors underwent not only neural tube differentiation, but also remained as mesenchymal cells in the tail bud (white arrowheads). A white small dot indicates the position of 31-33 somite level. White arrow is the boundary between the posterior end of epithelialized neural tube and mesenchymal SN precursors of the tail bud.

When these manipulated embryos were allowed to develop until HH17 (30ss), EGFP-positive cells were, as expected, found in the epithelialized neural tube posterior to somite level 31-33. Importantly, EGFP signals were also detected in the mesenchymal population in the tail bud (Fig. 2C, E; 6 out of 8). We further performed a serial transplantation to a third host embryo (Fig. 3F, G; n=6), and observed again EGFP-positive cells both in the secondary neural tube posterior to the 31-33 somite level and in the tail bud (4 out of 6). These results indicate the existence of self-renewing SN precursors in the tail bud.

**Figure 3.**
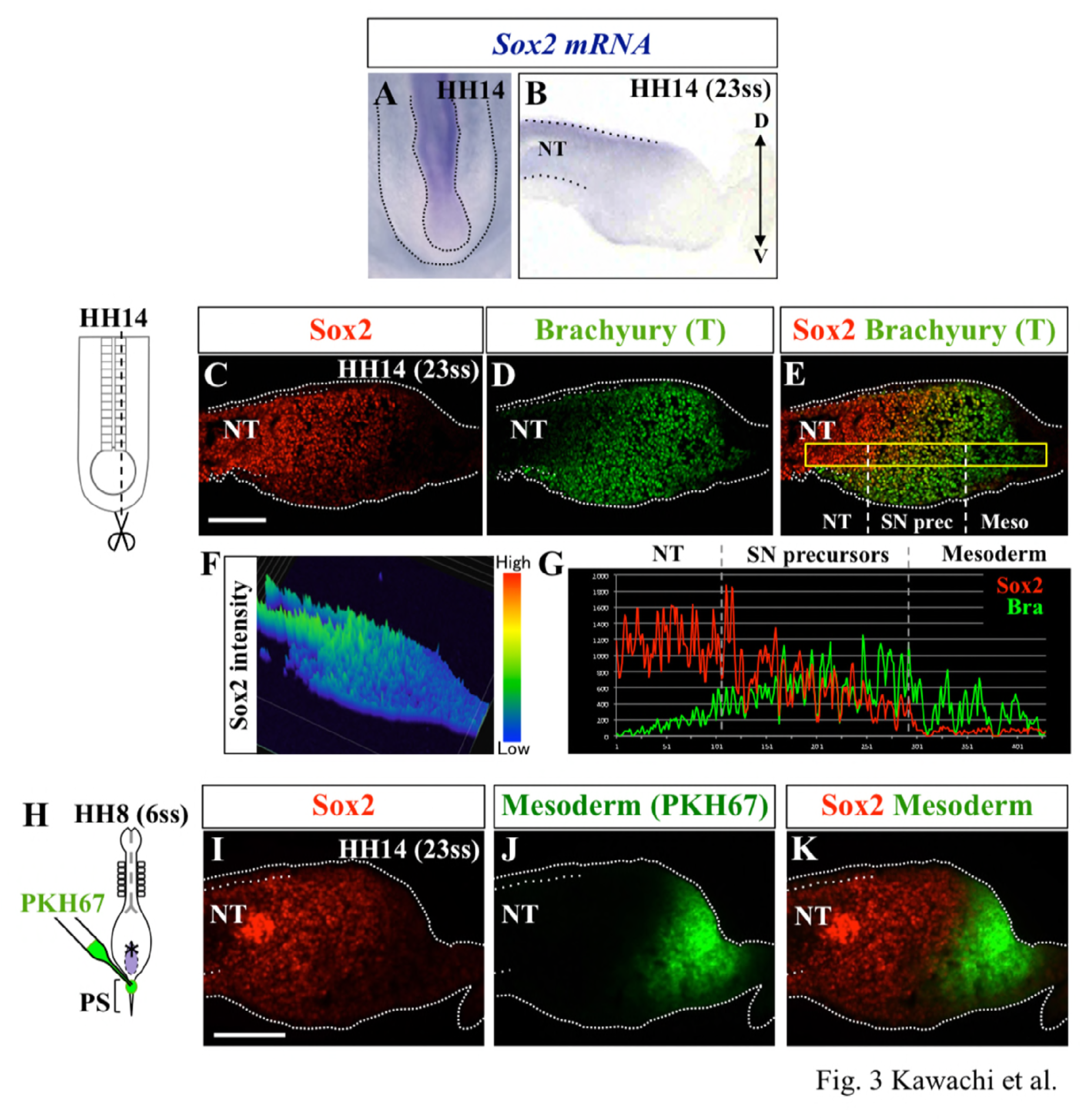
Graded expression levels of Sox2 in the SN precursor territory in the HH14 tail bud. (A, B) *In situ* hybridization for *Sox2* mRNA. Dorsal view (A) and sagittal section (B). (C-D) Immunohistochemistry for Sox2 and Brachyury (T) proteins in a para-sagittal section. (F, G) Graded expression of Sox2 was quantitatively shown, whereas the expression level of Brachyury was largely steady in the tail bud. (I-K) The Sox2-positive area overlaps with the SN-fated territory, but not with mesodermal cells derived from the anterior primitive streak (PKH67-labeled). Scale bar; 100 µm.

To more precisely locate these self-renewing cells in the SN precursor territory of the tail bud, we performed similar experiments, but this time we separated the anterior and posterior halves of the SN precursor domain, and transplanted each of them into its equivalent domain of stage-matched embryos (HH14). When the anterior half was transplanted, EGFP-positive cells were found solely in the differentiated neural tube posterior to the presumptive somite level 31-33 in HH17 embryos (Fig. 2J-L; 6 out of 7). In clear contrast, EGFP-cells from the posterior half of the SN precursor domain were found in both the epithelialized neural tube and undifferentiated mesenchymal population in the tail bud (Fig. 2M-O; 5 out of 7). Thus, SN precursors in the tail bud are subdivided at least into two different subpopulations: one is anteriorly located and specified to neural epithelialization/differentiation; and the other is the compartment of posterior cells maintained as undifferentiated. When the posterior half was transplanted, EGFP-positive cells in the secondary neural tube were located posteriorly to somite pair 35 and not to somite pairs 31-33, thereby suggesting that it takes 3-6 hrs for the self-renewing cells to be incorporated to the forming neural tube (one cycle of somite segmentation is 90 min in chickens) (Palmeirim *et al.*, 1997).

### Graded expression of Sox 2 in SN precursors

To explore the molecular mechanisms by which the two different subpopulations of SN precursors are regulated and distinguished, we paid attention to the expression of the *Sox2* gene, known to be active at the earliest stages of PN and SN neural tube formation (Uchikawa *et al.*, 2011). Indeed, the presence of *Sox2* mRNA is concomitant with neural epithelialization during the SN process (Shimokita & Takahashi, 2011). In the current study, we carefully examined expression levels of Sox 2 in the tail bud of HH14 (23ss) chicken embryos. In situ hybridization for *Sox2* mRNA revealed expression in the *anterior* domain of SN precursors, whereas signals were barely detectable in the *posterior* domain (Fig. 3A, B; n=6). We therefore switched to immunohistochemistry using anti-Sox2 antibody, allowing sensitive detection of the Sox2 protein. Confocal microscopic analyses virtualized nuclear Sox2 protein both in the anterior and posterior domains of SN precursors with graded signals deceasing posteriorly. This notion was supported by the quantitative evaluation (Fig. 3C, F; n=15).

We also compared the staining pattern of Sox2 with that of Brachyury (T) protein in the tail bud of HH14 embryos. It was previously shown that in very early mouse embryos, a specific epiblastic region that gives rise to both neural and mesodermal progenitors (NMPs) expresses both Sox2 and Brachyury (Bra), and since then, the Sox2^+^/Bra^+^ double-positive signal has been proposed as a marker for NMPs (Olivera-Martinez *et al.*, 2012, Wymeersch *et al.*, 2016, Gouti *et al.*, 2014). Contrasting with these proposals, we found that the Brachyury protein was widely distributed in the tail bud as shown in the sagittal section in Fig. 3D (n=10). Most importantly, the Bra^+^ region included the Sox2^+^ precursor territory, that is, almost the entire anterior half of the Bra^+^ domain was Sox2^+^/Bra^+^ double positive. Furthermore, unlike Sox2, the signal intensity of Bra was relatively uniform throughout in the tail bud, although it dropped abruptly following neural epithelialization, as expected (Fig. 3G; n=6). Thus, the mesenchymal cells of the HH14 tail bud were positive for Bra regardless of their developmental fate.

To further corroborate our observation that Sox2 expression was restricted to the neural fate in the tail bud, we double stained a sagittal section of HH14 tail bud for the Sox2 protein (red) and the fate of the anterior primitive streak-derived mesoderm (PKH-labeling). As expected, the mesoderm territory was negative for Sox2 (Fig. 3H; n=14).

These results revealed an intimate correlation between the differential levels Sox2 expression and different states of SN precursors in the tail bud: low and high levels of Sox 2 for self-renewing- and SN-specified populations, respectively.

### Augmented level of Sox 2 expression directed SN precursors to precocious differentiation/epithelialization

To know whether differential levels Sox2 expression are functionally related to the two groups SN precursors, we experimentally elevated the level of Sox2 expression in these cells. Similar to the experiments shown in Fig 1, the *in ovo* electroporation was carried out at HH8 (6ss) to target the preSN epiblastic region. However, if the overexpressed Sox 2 inhibited EMT/ingression of the epiblast, the electroporated cells would end up in neural epithelia at later stages, which might mislead interpretation. To avoid this possible confusion, we switched on the *Sox2* expression after the electroporated preSN-derived cells ingressed as mesenchyme, using the Tet-on expression system (Watanabe, et al., 2007).

As shown in Fig. 4A, three plasmids were co-electroporated: pCAGGS-mCherry (constitutively active), pBI-Sox2/EGFP (BI, previously called TRE, is a bidirectional promoter induced by doxycycline, an analog of tetracycline; Dox), and pCAGGS-Tet-On 3G (transcriptional activator which binds the tetracycline responsive element (TRE) only in the presence of Dox). When electroporated embryos reached HH14 (22ss), a Dox solution was administered. At the time of administration (0 hr), a decent amount of ingressed cells were already found in the tail bud, which expressed mCherry but not EGFP (Fig. 4A, B, E; n=7).

**Figure 4.**
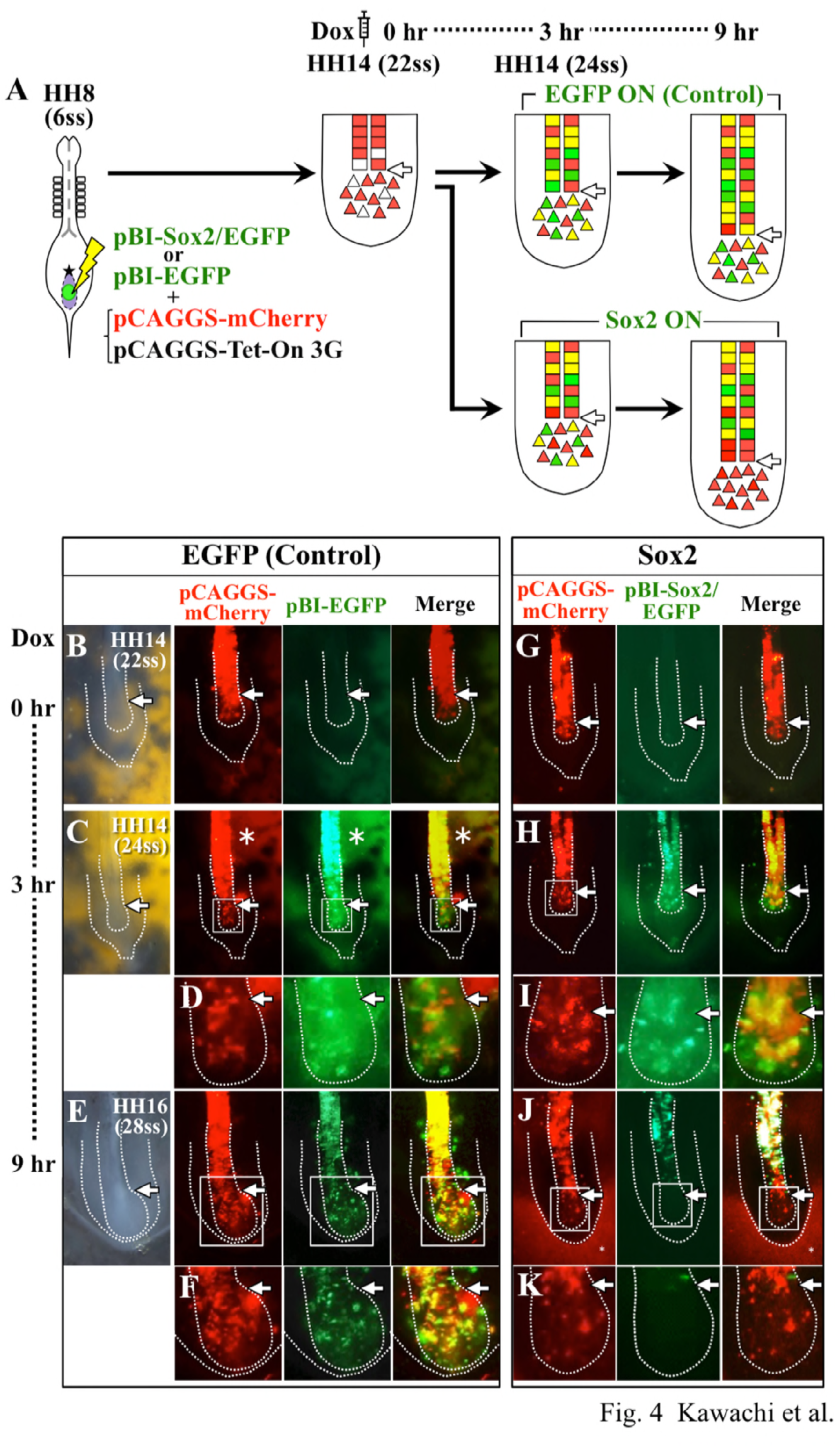
An experimentally augmented level of Sox2 caused precocious differentiation of SN precursors. (A) *Sox2* cDNA was overexpressed in a temporally controlled manner using the Tet-on system. The preSN region of HH8 embryos was electroporated with the plasmids indicated. When the embryo reached HH14, at which gene-electroporated cells emerged as mesenchyme in the forming tail bud, a Dox solution was administered (0hr). Note that at 3 hrs post-Dox in both control and Sox2-overexpressing embryos, three types of cells, green (EGFP), red (mCherry) and yellow (EGFP and mCherry) were observed in the tail bud (see text for details). However, at 9 hrs, Sox2-overexpressing cells were found solely in the formed neural tube, and not in the tail bud.

By 3 hrs post Dox at HH14 (24ss), EGFP/Sox2 started to be expressed in SN precursors in the tail bud, which is consistent with the previous report that a TRE-driven gene starts to be expressed after 3 hr of Dox injection (Watanabe, et al., 2007). It is known that the *in ovo* electroporation in chickens yields a mosaic pattern of transfected cells, and also that if two different plasmids, mCherry and EGFP, are co-electroporated, three types of transfected cells emerge: one double positive (mCherry^+^/EGFP^+^), and two single positives (mCherry^+^/EGFP^−^ or mCherry^−^/EGFP^+^). Indeed, at 3 hrs post Dox, these three types of cells were observed in *EGFP/Sox2*-electroporated SN precursors in the tail bud, which was comparable to EGFP-electroporated control cells. (Fig. 4A, C, F; n=7).

In contrast to 3 hrs, at 9 hrs post Dox, that is 6 hrs after the onset of *Sox 2* expression, distribution patterns of *EGFP/Sox2*-electroporated SN precursors were significantly different from those in control *EGFP*-electroporated embryos. In the control tail bud, all the three types, mCherry^+/^EGFP^+^, mCherry^+^/EGFP^−^ and mCherry^−^/EGFP^+^ cells, were observed (Fig. 4E, F; n=7). However, in *EGFP/Sox2*-electroporated embryos, only mCherry^+^ but no EGFP^+^ cells were detected in the tail bud (Fig. 4J, K; n=11). In addition, although the anteriorly formed neural tube contained double-positive mCherry^+/^EGFP^+^ cells, and single positive mCherry^−^/EGFP^+^ or mCherry^+^/EGFP^−^ cells, the nasant area of epithelialized cells at the boundary between the tail bud was preferentially occupied only by mCherry^+^/EGFP^−^ cells (Fig 4. J, K, near white arrow). Since we observed no sign of apoptosis in the tail bud in these embryos (not shown), it is most likely that the Sox2-elevated SN precursors underwent precocious differentiation/epithelialization.

### Inhibition of Sox2 prevented both differentiation/epithelialization and self-renewal ability of SN precursors

We further asked whether Sox2-deprived SN precursors would remain in a self-renewing state in the tail bud. To this end, we used Sox3HMG-EnR, known to repress endogenous Sox B type genes (Sox 1 to Sox 3) (Bylund *et al.*, 2003, Sasai, 2001). Since SN precursors in the tail bud express Sox2, but not Sox1 or Sox3 (Uchikawa *et al.*, 2011), electroporated Sox3HMG-EnR was considered to repress the activity of Sox 2. Using the Tet-on expression system shown in Fig. 4, *Sox3HMG-EnR* was Dox-induced to be expressed in a temporally controlled manner. This time, we electroporated pBI-Sox3HMG-EnR/ DsRed (which are driven bidirectionally by TRE), pCAGGS-EGFP, and pCAGGS-Tet-On3G. In addition, to precisely track the *Sox3HMG-EnR*-electroporated cells during secondary neural tube formation, we dissected gene-electroporated SN precursors from a tail bud of HH14 (22ss) embryo, and transplanted them homotopically into stage-matched non-electroporated embryos, into which Dox solution had been administered (Fig. 5A, B; n=5).

**Figure 5.**
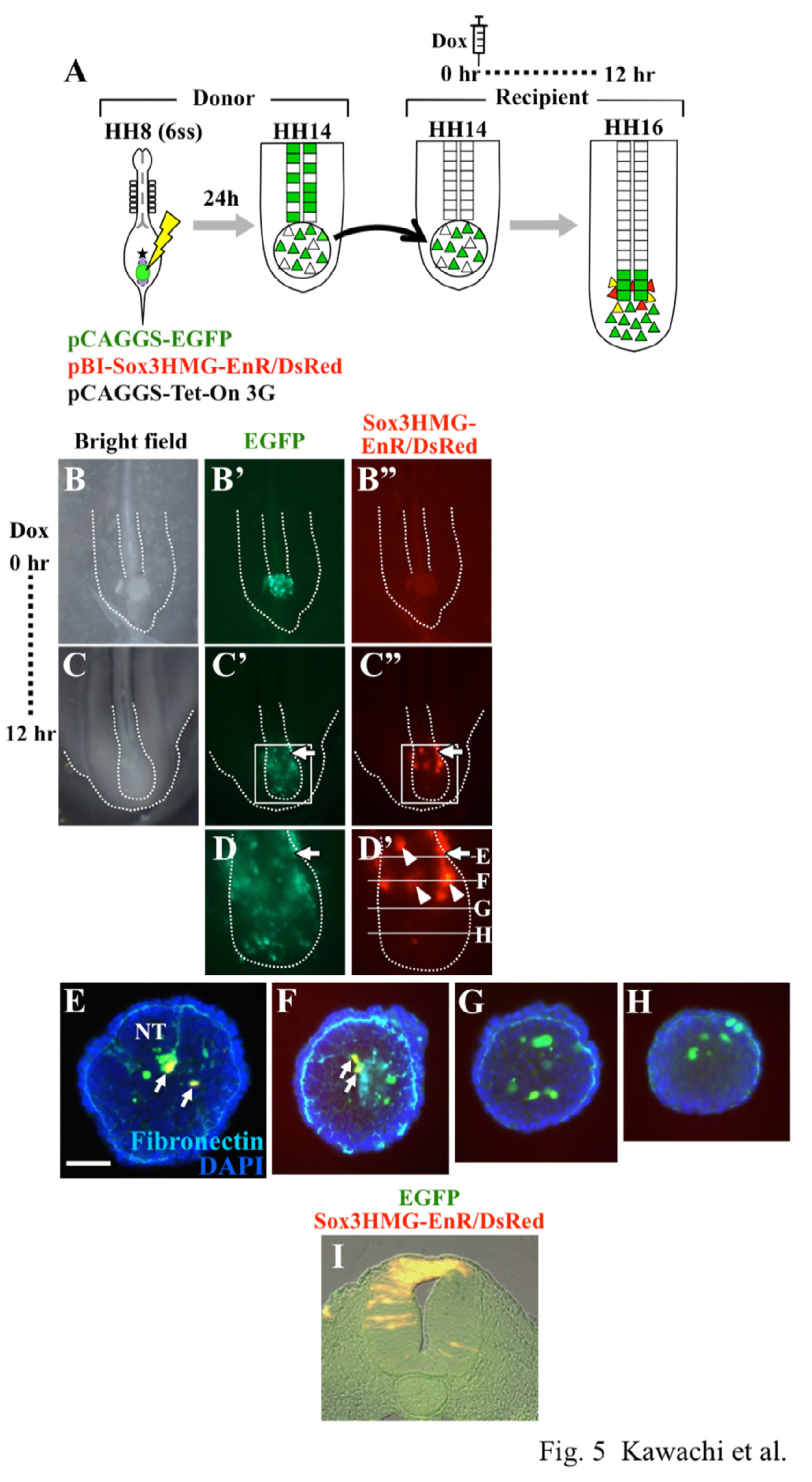
Sox2-deprived SN precursors failed to participate in the secondary neurulation. (A) *In ovo* electroporation and tet-on induced expression of *Sox3HMG-EnR* were carried out in a way similar to that shown in Fig. 4. In addition, Sox2-deprived SN precursors were precisely tracked following isotopic transplantation. (B) At the time of transplantation (0 hr), no signal for the expression of Sox3HMG-EnR was seen yet. (C) By 12 hrs post Dox, *Sox3HMG-EnR*-expressing SN precursors cells (red), whose number was smaller than that of control (EGFP) cells, failed to be incorporated into the secondary neural tube formation (E, F). (G, H) In the posterior region of the SN precursor territory, no Sox3HMG-EnR-expressing cells were observed. Scale bar; 50 µm.

By 12 hrs post Dox, when embryos reached HH16, DsRed (Sox3HMG-EnR) signals were detected (the onset of DsRed detection after Dox was delayed compared to mCherry due to the protein configuration) (Fig. 5A, C; n=5). Notably, at this stage, only a few DsRed^+^ cells were recognized in the tail bud probably due to the cell death of *Sox3HMG-EnR*-expressing cells, and even this remnant of remaining cells failed to participate in the secondary neural tube (Fig. 5C, D; n=5). It clearly contrasts with the observations shown in Fig 2, where transplanted EGFP^+^ (normal) cells were actively incorporated into the secondary neural tube. It is unlikely that Sox2-inhibited cells were converted to mesoderm. In transverse sections, Sox 2-inhibited cells (red) in the anterior domain of SN precursors, which would normally express a high level of Sox 2, were located near the neural tube within the territory of SN precursors (Fig. 5E, F; n=4). These sections also confirmed that the posterior domain of SN precursors, which would normally be of low Sox 2, did not contain Sox 2-inhibited cells (Fig. 5D’, G, H; n=4). The distribution of single positive EGFP^+^ cells, which did not receive the *Sox3HMG-EnR* gene, was comparable to normal cells (Fig. 5D compare to Fig. 2; n=4). Collectively, these results suggest that a high level of Sox2 is required for the anterior SN precursors to correctly be incorporated into the secondary neural tube. And even the low level of Sox 2 activity in the posterior SN precursors is indispensable, particularly for the self-renewing cells to survive.

The loss of *Sox3HMG-EnR*-electroporated cells was specific to SN precursors since when the expression of this gene was Dox-induced in already epithelialized cells in the secondary neural tube, no detectable effects were seen until HH16 (Fig. 5I; n=7).

## Discussion

Using the direct labeling of SN precursors, SN-specific transplantations, and SN-specific gene manipulations, we have uncovered cellular and molecular mechanisms underlying the SN-mediated neural elongation in the posterior region of chicken embryos. SN precursors occupy the anterior half of the tail bud at HH14, whereas the posterior half is populated by paraxial mesodermal precursors. The SN precursor territory was further divided into two distinct regions; one is its posterior half with self-renewing cells, and the other is the anterior half more specified to the neural tubulogenesis (Fig. 6). Sox2 plays an important role in the regulation of these two distinct states of cells: high and low levels of Sox2 activities endow the neural-specified and self-renewing states, respectively.

**Figure 6.**
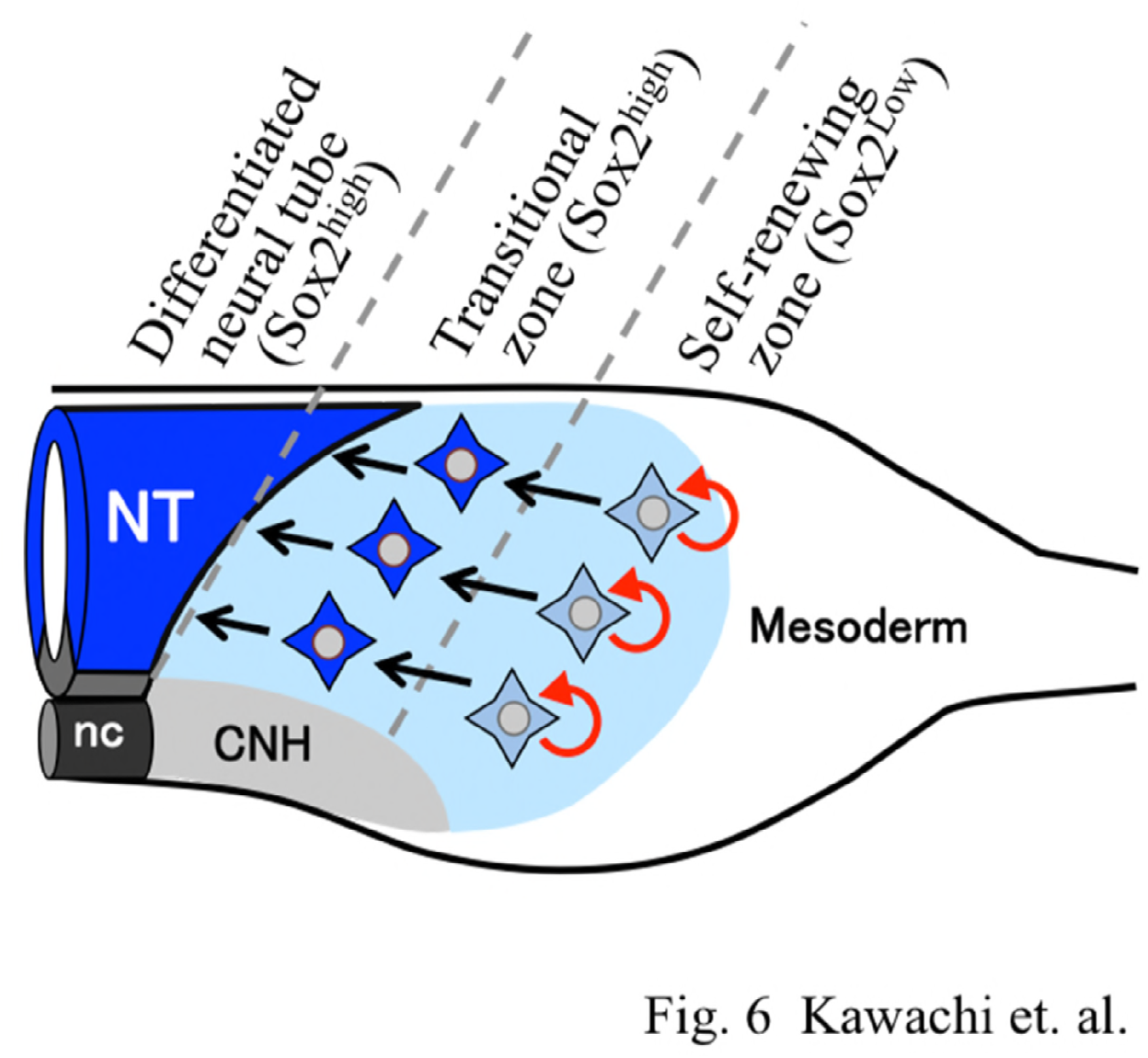
A summary diagram of the SN precursors in the tail bud of chicken embryos. SN-fated cells populate the anterior half of the mesenchymal mass of the tail bud, whereas the rest of the tail bud is populated by mesodermal precursors (paraxial mesoderm) and chord neural hinge (CNH). The SN precursor territory (light blue) is further divided into two subpopulations: self-renewing cells posteriorly and more neural-specified transitional cells anteriorly. Sox 2 displays posteriorly decreasing levels of expression in the SN precursor territory. High and low levels of Sox 2 are important for neural differentiation/epithelialization of transitional zone, and survival of self-renewing cells, respectively.

### A discrete territory of SN precursors in the tail bud

The tail bud has long been appreciated to be critical for the posterior body elongation, and the existence of precursors has been shown that give rise to the secondary neural tube, and/or paraxial mesoderm, and the chord neural hinge (CNH), although their precise positions in the tail bud have been under debates (Cambray & Wilson, 2002, Catala *et al.*, 1996, Catala *et al.*, 1995, McGrew *et al.*, 2008, Selleck & Stern, 1991, Olivera-Martinez *et al.*, 2012, Tucker & Slack, 1995). Recent advances in genetic labeling of earlier epiblast cells in mice proposed that neuromesodermal common progenitors (NMPs) exist not only in the early epiblast but also in the tail bud (Tzouanacou *et al.*, 2009, Wymeersch *et al.*, 2016). However, no direct fate mapping of the tail bud-constituting cells has been conducted in amniotes.

In this study, we have extended our previous findings that the SN precursors can directly be traced in the tail bud (Shimokita and Takahashi, 2011). These SN precursors derive from the preSN region in the epiblast at HH8, which is located in between Hensen’s node and the anterior end of the primitive streak (Fig. 1). By focusing on the tail bud at HH14 (24ss), which displays a representative feature of this structure, we have delineated SN- and mesoderm-harboring regions. The SN precursors occupy the anterior half, whereas the posterior half is populated by mesodermal cells derived from the anterior primitive streak, leaving the CNH unlabeled (Fig. 1J). This delineation is striking to us because several previous studies of axial elongation postulated prominent NMPs in the tail bud, although these studies traced more naïve cells residing in the epiblast at earlier stages than those used in the current study (Selleck & Stern, 1991, Wymeersch *et al.*, 2016, Taniguchi *et al.*, 2017). Thus, our study clarified, for the first time, different fates of mesenchymal cell populations in the tail bud, at least at the HH14 (24ss) stage. Our results do not exclude the existence of NMPs in the tail bud, since preSN-labeling rarely yields cells located in the medial edge of presomitic mesoderm (see Fig. 4E). Nevertheless, a vast majority of the tail bud cells are differentially fated to either SN, mesoderm, or CNH.

### Posteriorly positioned SN precursors possess self-renewing ability

The SN precursors are constantly present in the tail bud despite their continuous participation in the secondary neural tube formation, and we had reasoned that SN precursors in the tail bud might contain stem (-like) cells. Specific tracing of the SN precursors in the tail bud has facilitated to test this, and we have indeed shown that such stem cells exist in the tail bud by a serial transplantation of EGFP-labeled SN precursors (Fig. 2A). Furthermore, these self-renewing cells are preferentially located in the posterior half of the SN precursor territory (Fig. 2, Fig. 6). These posterior cells undergo both self-renewal and its subsequent neural tubulogenesis (epithelialization), whereas the anterior half cells only participate in the tubulogenesis without staying in the tail bud. Thus, the SN precursors contain at least two subpopulations, one is posteriorly positioned self-renewing cells, and the other is anteriorly located cells in a transitional state (Fig. 6). This is the first identification of SN stem cell population in the tail bud. Whether a single stem cell generates both self- and differentiating cells awaits further analyses.

### AP gradient of Sox2 expression regulates the binary decision between self-renewing and transitional states of SN precursors

Sox2 displays a posteriorly decreasing gradient of expression within the SN precursor territory in the tail bud, which contrasts with a relatively steady level of Brachyury (T) expression both in the SN- and mesodermal precursors (Fig. 3). We have provided evidence that the AP gradient of Sox2 is important for the binary decision between the self-renewing and transitional states of SN precursors. With gain-of-function experiments, we have found that a high level of Sox2 directs cells to the differentiation/epithelialization of the secondary neural tube. Loss-of-function experiments have unexpectedly shown that even a low level of Sox2 is important for the posterior SN precursors. Sox2 has been implicated in normal neural development, pluripotency of stem cells, and survival of neural progenitors (Feng *et al.*, 2013, Hagey & Muhr, 2014, Hutton & Pevny, 2011, Kondoh & Lovell-Badge, 2016). In particular, a low level of Sox2 is important for the proliferation of neural stem cells in adult brain (Hagey & Muhr, 2014). It is possible that the self-renewing SN precursors identified in the current study might serve as a novel source for therapeutic treatments of neural diseases.

It is yet to be determined what generates the AP gradient of Sox 2 expression in the SN precursor territory. It is known that the preSN region in the epiblast of HH8 embryo, from which SN precursors arise, is devoid of Sox2, although its adjacent neural plate/epiblast (i.e. primary neurulation tissues) expresses this gene, and also that this preSN-specific inactivation of Sox2 requires BMP4 (Takemoto *et al.*, 2006). It is conceivable that the AP gradient of Sox 2 expression in the tail bud is also regulated by BMPs expressed in the tail bud and its surrounding tissues (Takemoto *et al.*, 2006).

Previous studies of axial elongation have regarded Sox2^+^/Bra^+^ double positive signal as a marker for an NMPs cell population in the early epiblast (Olivera-Martinez *et al.*, 2012, Wymeersch *et al.*, 2016, Delfino-Machin *et al.*, 2005, Tsakiridis *et al.*, 2014, Yoshida *et al.*, 2014). However, in the current study, we have explicitly demonstrated that a majority of the SN precursors in the tail bud are Sox2^+^/Bra^+^ double-positive, even though they are *not* NMPs. Thus, the fate of Sox2^+^/Bra^+^ double positive cells must vary according to the developmental context. It is also possible that Sox2^+^/Bra^+^ cells might have high potential to easily change their fate upon encountering external signals. We are currently investigating differentiation potential of SN- and paraxial mesodermal precursors in the tail bud based on the developmental lineages delineated in the current study.

Lastly, it is of interest speculating that the self-renewing ability of the SN precursors might reflect the stemness of adjacent mesoderm (Iimura & Pourquie, 2006). When mesodermal cells that are derived from the anterior primitive streak of HH4-6 embryos give rise to the medial half of somites, they do so via the tail bud where some of these cells stay for a long period of time and serve as self-renewing cells (Iimura & Pourquie, 2006). According to our findings, such mesodermal stem cells should be those located in the posterior half of the tail bud at HH14, and more ventrally at HH17 (Fig. 1). In this way, both the mesodermal stem cells contributing to the medial half of somites and the SN stem cells found in this study are possibly adjacent to each other in the tail bud (Fig. 6). It is plausible that these differentially fated but similarly self-renewing cells would share signals for the endowment of the stemness. Whether these signals include Fgfs and Wnts, which are predominantly expressed in the posterior region of the elongating body awaits further analyses.

## Acknowledgements

We thank Dr. Scott F. Gilbert for helpful discussion and careful reading of the manuscript. We thank Drs. Y. Sasai (RIKEN, deceased) and K. Fukuda (Tokyo Metropolitan University) for kind gifts of Sox3HMG-EnR and Sox 2, respectively. This work was supported by JSPS KAKENHI: Grant-in-Aid for Scientific Research (B), SPIRITS (Kyoto University), and AMED (JP17gm0610015). T.K. and E. S. were fellows of JSPS (DC1).

